# Autophagic signaling promotes systems-wide remodeling in skeletal muscle upon oncometabolic stress

**DOI:** 10.1101/2020.10.13.338202

**Authors:** Anja Karlstaedt, Heidi Vitrac, Rebecca L. Salazar, Benjamin D. Gould, Daniel Soetkamp, Weston Spivia, Koen Raedschelders, An Q. Dinh, Anna G. Guzman, Lin Tan, Stavros Azinas, David J.R. Taylor, Walter Schiffer, Daniel McNavish, Helen B. Burks, Roberta A. Gottlieb, Philip L. Lorenzi, Blake M. Hanson, Jennifer E. Van Eyk, Heinrich Taegtmeyer

## Abstract

About 20-30% of cancer-associated deaths are due to complications from cachexia which is characterized by skeletal muscle atrophy. Metabolic reprogramming in cancer cells causes body-wide metabolic and proteomic remodeling, which remain poorly understood. Here, we present evidence that the oncometabolite D-2-hydroxylgutarate (D2-HG) impairs NAD^+^ redox homeostasis in skeletal myotubes, causing atrophy via deacetylation of LC3-II by the nuclear deacetylase Sirt1. Overexpression of p300 or silencing of Sirt1 abrogate its interaction with LC3, and subsequently reduced levels of LC3 lipidation. Using RNA-sequencing and mass spectrometry-based metabolomics and proteomics, we demonstrate that prolonged treatment with the oncometabolite D2-HG in mice promotes cachexia *in vivo* and increases the abundance of proteins and metabolites, which are involved in energy substrate metabolism, chromatin acetylation and autophagy regulation. We further show that D2-HG promotes a sex-dependent adaptation in skeletal muscle using network modeling and machine learning algorithms. Our multi-omics approach exposes new metabolic vulnerabilities in response to D2-HG in skeletal muscle and provides a conceptual framework for identifying therapeutic targets in cachexia.

## Introduction

Skeletal muscle wasting is a hallmark of cachexia, which is associated with systemic multi-organ diseases like heart failure and cancer (Evans *et al*, 2008; Fearon *et al*, 2011). Patients with cachexia have a poor prognosis in advanced disease stages, and about 20 to 30% of cancer-associated deaths are due to cachexia (Fearon *et al.*, 2011). The severity of skeletal muscle atrophy can be associated with a specific tumor type rather than with tumor size and burden (Fearon *et al.*, 2011). Severe muscle loss is commonly associated with hematological malignancies (e.g., leukemia), pancreatic tumors, non-small cell lung tumors, and gastrointestinal tract tumors (Lok, 2015). The genetic and metabolic basis for the development of skeletal muscle atrophy is elusive and limits our ability to develop targeted therapeutic strategies without those side effects.

Recent studies indicate that skeletal muscle atrophy during cachexia is characterized by increased proteome proteolysis (Gallagher *et al*, 2012; Salazar-Degracia *et al*, 2019), impaired mitochondrial metabolism (Friesen *et al*, 2015), inefficient ATP provision (VanderVeen *et al*, 2017), dysfunction of the electron transport chain (VanderVeen *et al.*, 2017), and extensive cellular lipid remodeling (Kir & Spiegelman, 2016). These molecular changes are not just a severe complication of tumor growth, but rather a consequence of the metabolic reprogramming driven by the biology of tumors (Gallagher *et al.*, 2012). The metabolic phenotype of tumors is often characterized by the accumulation of specific intermediates or oncometabolites. The oncometabolite D-2-hydroxyglutarate (D2-HG) promotes contractile dysfunction in the heart (Karlstaedt *et al*, 2016). Mutations of isocitrate dehydrogenase (IDH) 1 and 2 lead to the gain of a neo-morphic enzymatic function and production of the oncometabolite D2-HG, which accumulates to millimolar level in both tumors and the bloodstream (DiNardo *et al*, 2013; Kranendijk *et al*, 2010a; Kranendijk *et al*, 2012; Kranendijk *et al*, 2010b). IDH 1 and 2 mutations are observed in several types of cancers including glioblastomas (Louis *et al*, 2016; Miller *et al*, 2019), acute myeloid leukemia (AML) (Cancer Genome Atlas Research *et al*, 2013), chondrosarcoma (Lu *et al*, 2013), and cholangiocarcinoma (Amary *et al*, 2011; Pansuriya *et al*, 2011). D2-HG suppressed energy-provision, and impaired NADH regeneration in the heart through inhibition of α-ketoglutarate dehydrogenase (α-KGDH) (Karlstaedt *et al.*, 2016). Prolonged exposure to D2-HG was associated with both heart and skeletal muscle atrophy in mice, which prompted the hypothesis that oncometabolic stress mediates autophagy or proteasomal degradation of muscle proteins. Protein synthesis and breakdown in skeletal muscle is regulated by a variety of factors, including nutrient supply, hormone concentrations, and physical activity. Muscle mass homeostasis is maintained through balancing rates of protein synthesis and degradation, the latter including the ubiquitin/proteasome pathway (UPP), autophagy, caspases, cathepsins, and calcium-dependent calpains. Previous studies have demonstrated the involvement of the UPP with increased mRNA expression of muscle-specific E3 ubiquitin ligases, including atrogin-1 or maf box 1 (MafBx1) and muscle-specific ring finger protein 1 (Murf1 or Trim63), during the acute stress response in skeletal muscle (Bodine *et al*, 2001; Gomes *et al*, 2001). Several studies have implicated both protein synthesis and degradation pathways in animal models of cancer-associated muscle wasting (Gallagher *et al.*, 2012; Smith & Tisdale, 1993). Similarly, increased α-KG metabolism prevents activation of autophagy (Baracco *et al*, 2019).

Our goal was to characterize the metabolic, proteomic, and genomic changes in skeletal muscle that occur in response to D2-HG and understand which biological processes are involved in driving adaptation. We devised a multi-omics approach to study the system-wide impact of D2-HG on skeletal muscle metabolism and physiology in cell culture and mouse models. We link gene and protein expression to metabolic changes and reveal the complex interplay between biological processes in response to acute and chronic stress. We show that the NAD^+^-dependent sirtuin 1 signaling pathway integrates protein and nutrient signals to regulate autophagy in response to oncometabolic stress. Targeted metabolomics revealed that changes in autophagy activation affect the metabolism of energy-providing substrates in skeletal muscle cells. Importantly, using an integrative multiomics approach, we show that these changes induce a system-wide adaptive response in mouse models that affect both gene and protein expression profiles. This study highlights the importance of system-wide profiling to decipher complex biological adaptation in response to oncometabolic stress and provides an integrated approach to modeling cancer cachexia in the context of IDH-mutant cancers.

## Results

### Inhibition of α-KGDH causes metabolic remodeling in skeletal muscle cells

The oncometabolite D2-HG inhibits α-KG-dependent enzymes, including α-KGDH, leading to decreased Krebs cycle flux and impaired energy provision in the heart (Karlstaedt *et al.*, 2016). We previously showed that prolonged treatment with D2-HG is associated with heart and skeletal muscle atrophy (Karlstaedt *et al.*, 2016). To identify metabolic pathways that are driving skeletal muscle adaptation in response to oncometabolic stress, we applied targeted metabolomic analyses using liquid chromatography and mass spectrometry (LC-MS/MS) on L6 myotubes. These myotubes are derived from rat L6 myocytes (L6Ms, rat skeletal muscle cell line) and consist of fused myoblasts, which retain the ability to contract (Oberg *et al*, 2011). We cultured L6 myotubes with either phosphate buffered saline (PBS, control), D2-HG (1.0 mmol/L) or dimethyl alpha-ketoglutarate (DMKG, 1.0 mmol/L) for 24 h in defined nutrient-rich media (**Figure 1A**). DMKG is a membrane-permeable ester of α-KG that is cleaved to α-KG in the cytoplasm, consequently increasing α-KG levels (Shah *et al*, 2010). We found that the both D2-HG and DMKG treatment decreased the availability of critical metabolic intermediates (e.g., glucose 6-phosphate, sedoheptulose-1,7-bisphosphatase) (**Figure 1B** and **C**). D2-HG treatment reduced NADH and increased the overall NAD^+^/NADH redox ratio in L6 myotubes, whereas treatment with DMKG did not affect these metabolites (**Figure 1C**), indicating that these changes are purely D2-HG-dependent. We integrated the targeted LC-MS/MS metabolomic data into pathway enrichment analysis to assess metabolic remodeling based on pathways and chemical similarity using MetaMapp (Barupal *et al*, 2012) and Cytoscape (Shannon *et al*, 2003) (**Figure 1D**). The resulting network is composed of 101 metabolites with 615 metabolite-metabolite interactions (MMIs). We identified enrichment of metabolites in redox homeostasis (e.g., NADH, pentose phosphate pathway), and amino acid and glucose metabolism. These findings are consistent with previous studies in heart tissue suggesting that cancer cells producing D2-HG have a systemic metabolic impact on muscle tissue (Karlstaedt *et al.*, 2016).

**Figure 1.**
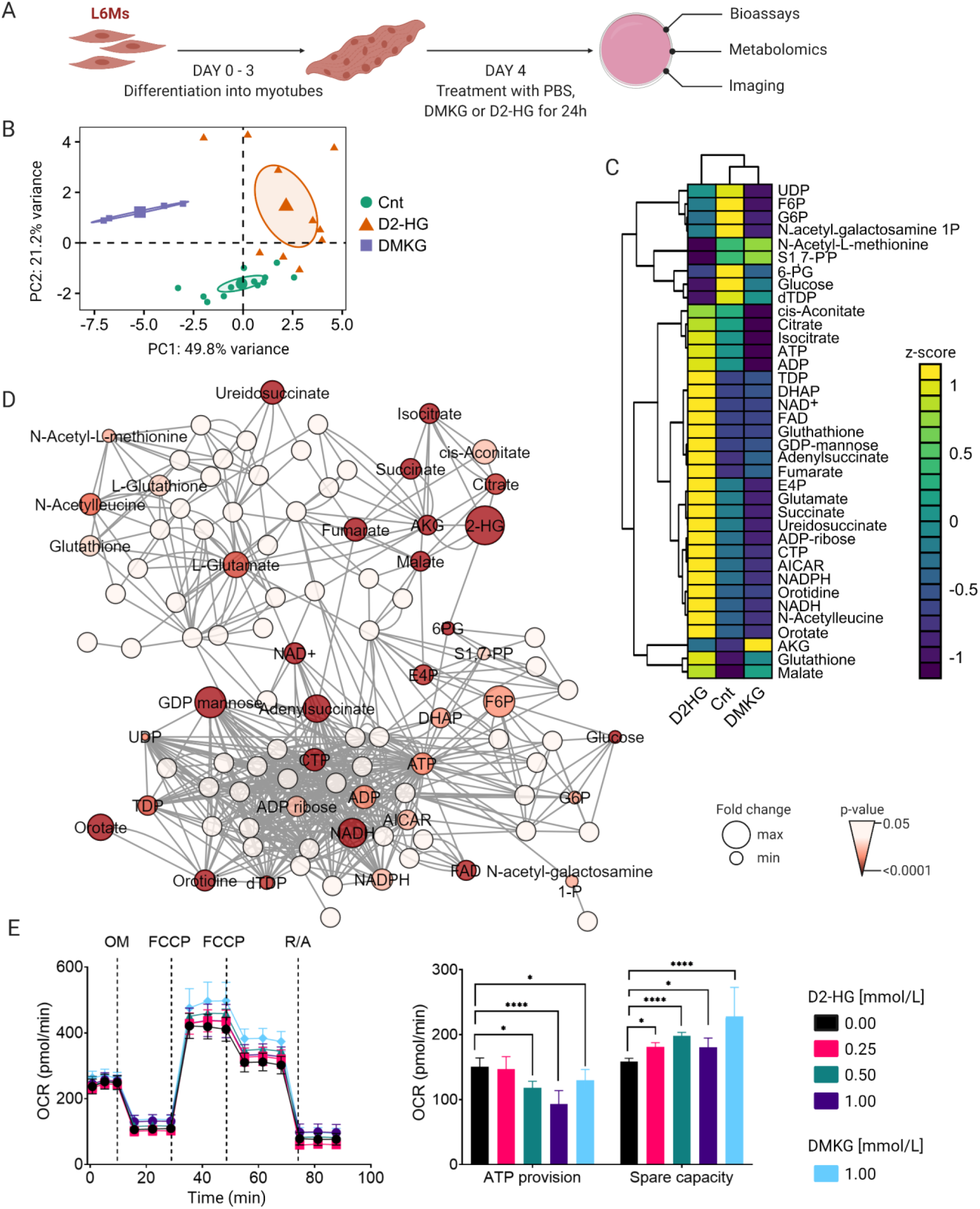
D2-HG causes metabolic remodeling in differentiated L6Ms. (**A**) Schematic of *in vitro* experimental workflow and protocol. (**B**) PCA of targeted metabolomics using LC-MS/MS in L6 myotubes treated with phosphate-buffered saline (Cnt), D2-HG (1 mmol/L), or DMKG (1 mmol/L) for 24 h. Each data point represents a biological replicate. Data is log2 normalized. (**C**) Heatmap showing unsupervised hierarchical clustering of metabolites that are altered (*P*-value<0.05, FDR=10%) in the metabolomics dataset used in (B). The color-coded z-score illustrates the increase or decrease in metabolite concentrations upon D2-HG treatment. (**D**) Network visualization of targeted MS-based metabolomics data. Nodes represent metabolites and edges represent metabolite-metabolite interactions. Nodes are color-coded by *P-*value and their size represent the median fold change relative to untreated sample. (**E**) Oxygen consumption rate (OCR) of L6 myotubes in response to D2-HG (0 to 1 mmol/L) was determined following sequential addition of oligomycin (OM, 1 μmol/L) to measure ATP-linked OCR, Carbonyl cyanide 4-(trifluoromethoxy)phenylhydrazone (FCCP, 0.2 μmol/L) to determine maximal respiration, and rotenone and antimycin (R/A, 1 μmol/L each) to determine the non-mitochondrial respiration as indicated. ATP-linked OCR and spare capacity were determined in L6 myotubes treated with the indicated dose of D2-HG (0 to 1 mmol/L) (*n* = 7 technical and biological replicates per group). Data are mean ± s.d. \**P*-value<0.05, \*\*\*\**P*-value<0.0001. Statistical analysis using two-way ANOVA with post-hoc Tukey’s multiple comparisons test.

NAD^+^ is a critical co-factor and reducing equivalent for a variety of metabolic processes. Alteration in the NAD^+^/NADH redox potential can directly affect the function of enzymatic reactions and indicate functional impairment of mitochondria. Thus, we reasoned that if α-KG levels are elevated in L6 myotubes as a response to impaired Krebs cycle flux, the increase in NAD^+^ levels might be accompanied by a decreased mitochondrial function and ATP provision. Using Agilent Seahorse XF Cell Mito Stress Tests, we confirmed that both D2-HG and DKMG treatment decreases the ATP-linked oxygen consumption rate (OCR) and increases the mitochondrial spare capacity in L6 myotubes in a concentration-dependent manner (**Figure 1E**, **Supplementary Figure 1A**). Intriguingly, we did not observe changes in mitochondrial function or ATP-related OCR at D2-HG concentration between 0.06 and 0.125 mmol/L (**Supplementary Figure 1A**). These observations are consistent with previous reports that D2-HG or L2-HG may accumulate under physiologic conditions without a notable effect on enzymatic activities in certain cancer cell lines and tumors (Intlekofer *et al*, 2015; Intlekofer *et al*, 2017; Nadtochiy *et al*, 2015). Together, these data suggest that moderately high level of D2-HG cause mitochondrial dysfunction in L6 myotubes, which parallels broad metabolic remodeling through increased amino acid metabolism and increased demand for glutamine.

### Deacetylation of LC3 is driving autophagy activation

Recent studies indicate that increased α-KG levels, and disrupted NAD+ redox homeostasis and ATP provision during nutrient-starvation induce autophagic flux and proteasomal degradation (Baracco *et al.*, 2019; Marino *et al*, 2014b). Thus, we reasoned that the observed metabolic changes in D2-HG-treated L6 myotubes might be accompanied by an acute increase in protein degradation pathways. To explore this possibility, we assessed the initiation and flux of autophagosome formation in L6 myotubes through quantification of microtubule-associated protein 1 light chain 3-II (LC3-II) using western blotting. L6 myotubes were cultured in defined nutrient-rich media and treated with or without D2-HG (0.5 and 1.0 mmol/L) in presence of bafilomycin A1 (BafA1). BafA1 inhibits the maturation of autophagosomes by blocking the fusion between autophagosomes and lysosomes, thus preventing LC3-II degradation and allowing to measure autophagic flux (Yamamoto *et al*, 1998). We found that D2-HG increases the lipidation of LC3 with and without bafilomycin A1 (BafA1, 200 nmol/L) (**Supplementary Figure 1B**). In contrast, the supplementation of DMKG maintained low autophagy levels (LC3-II and p62 expression) both with and without BafA1 conditions *in vitro* (**Supplementary Figure 1C**). The fusion between autophagosomes and lysosomes can be visualized using L6Ms expressing green fluorescent protein (GFP) fused with LC3 and LysoTracker (see **Methods** for details) as markers for autophagosomes and lysosomes. In untreated L6Ms (PBS; control), GFP-LC3 puncta were diffusely localized in the nucleus and cytosol (**Supplementary Figure 2A**). BafA1 treatment alone caused a significant increase in GFP-LC3 puncta formation in the cytosol and autolysosome formation in L6Ms, reflecting the basal level of autophagy activation (**Supplementary Figure 2A**). In contrast, D2-HG treatment increased the number of cytoplasmic puncta and the majority of GFP-LC3 colocalized with LysoTracker consistent with increased autophagic flux (**Supplementary Figure 2A**). We also measured the expression of critical genes involved in proteasomal degradation and autophagy: Beclin1, LC3, MafBx1, Murf1, and protein 62 (p62) were measured over 24 hrs. Gene expression of both LC3 and Murf1 increased within 12 h of treatment *in vitro*, reaching a plateau after 16 h and staying significantly increased at 24 h (**Supplementary Figure 2B** and **C**). In contrast, MafBx1 mRNA levels remained unchanged between experimental groups during the 24 h experimental protocol (**Supplementary Figure 2D**). Beclin1 mRNA levels increased after 2 h in D2-HG-treated cells (**Supplementary Figure 2E**), while p62 expression decreased after 8 h before returning to the baseline level (**Supplementary Figure 2F**). These results demonstrate that D2-HG increases autophagic flux and proteasomal degradation of proteins in skeletal muscle cells.

Mitochondrial function is directly linked to ATP provision and NAD-redox homeostasis. We observed both impaired ATP provision and increased NAD^+^ levels in response to oncometabolic stress. The AMP-activated protein kinase (AMPK) and mammalian target of rapamycin (mTOR) are crucial cellular energy and growth sensor proteins regulated by ATP levels in the cell. The total and phosphorylated protein expression of AMPK and mTOR were not increased after treatment with D2-HG or BafA1 (*P*-value>0.05 for all groups; **Supplementary Figure 3A** and **B**), suggesting that atrophy is activated by other mechanisms in our model. A second possible mechanism is the post-translational modification of proteins in response to metabolic stress. The NAD^+^-dependent deacetylase sirtuin-1 (Sirt1) promotes autophagy through de-acetylation of LC3 lysine (**Figure 2A**). In L6 myotubes, D2-HG treatment increases the co-localization of LC3 and Sirt1, which, in turn, decreases the acetylation of LC3 (**Figure 2B**). We next developed single and double mutants of LC3 with replacement of lysine at positions 49 and 51 by either arginine (LC3-K49R, LC3-K51R and LC3-K49R-K51R) or glutamine (LC3-K49Q, LC3-K51Q and LC3-K49Q-K51Q). These mutations allowed to mimic a decrease (K to R) or an increase (K to Q) in the acetylation level of LC3 (Huang *et al*, 2015). GFP-tagged single and double mutants were transfected into L6Ms and further differentiated into myotubes to assess their acetylation after treatment with or without D2-HG (1.0 mmol/L) for 24 h. Lysine-to-arginine single replacement at position 49 or 51 reduced LC3 acetylation, and the double mutant (LC3-K49R-K51R) showed the lowest acetylation (**Figure 2C**). Upon treatment with D2-HG, we detected increased deacetylation of LC3 in L6 myotubes expressing LC3-K49R or LC3-K51R. LC3 acetylation was almost completely abolished in LC3-K49R-K51R mutants and associated with increased LC3 lipidation (**Figure 2C**). Conversely, we found increased acetylation of LC3 in L6 myotubes expressing GFP-tagged LC3-K to Q mutants treated with and without D2-HG (**Supplementary Figure 3C** and **3D**). In all mutants, LC3 acetylation was decreased in D2-HG cells compared to untreated cells, indicating that the oncometabolite D2-HG increases the flux of deacetylation *in vitro* (**Supplementary Figure 3C**). We then measured the formation of GFP-LC3 containing puncta in GFP-tagged LC3-KR mutants through live-cell fluorescent microscopy. Untreated L6 myotubes expressing LC3-K48R, LC3-K51R or LC3-K48R-K51R showed a nuclear and cytoplasmic distribution like wild type (WT) LC3, which was also reflected in the amount of GFP-LC3 containing puncta per cell (**Figure 2D**). In contrast, treatment with D2-HG caused 2-to 3-fold increased formation of GFP-LC3 puncta in the cytosol (**Figure 2D**). Our findings indicate that the deacetylation of LC3 is the primary driver of autophagy activation in D2-HG treated L6 myotubes. Next, we tested whether the activation of autophagy is attenuated by increasing the deacetylation of LC3. We used two strategies to test this hypothesis. First, we silenced the expression of Sirt1 using siRNA and non-targeting negative controls to assess whether Sirt1 is solely driving the lipidation of LC3. Secondly, we overexpressed the acetyltransferase p300 in L6 myotubes using plasmid to test whether increased protein acetylation would counteract the effect of D2-HG. Previous studies have shown that p300 is a primary regulator of autophagy through protein acetylation during nutrient limitation (Lee & Finkel, 2009). At baseline, protein expression of both Sirt1 and p300 were increased in response to D2-HG **(Figure 2E**). Silencing Sirt1 attenuated both the total LC3-II level and the LC3-II to LC3-I ratio compared to WT conditions within 24 h (**Figure 2F**). Nonetheless, both total LC3-II level (*P*-value = 0.0004; q-value = 0.0008) and LC3-II to LC3-I ratio (P-value = 0.02; q-value = 0.023) were still significantly increased compared to WT conditions. These data suggest that other pathways may initiate LC3 lipidation in the presence of D2-HG. In contrast, overexpressing p300 **(Figure 2G**) fully attenuated total LC3-II levels (*P*-value = 0.5; q-value = 0.7) and LC3-II to LC3-I ratios (*P*-value = 0.6; q-value = 0.7) within 24 h, thus counteracting the effect of D2-HG. Our data indicate that NAD^+^ redox changes and the deacetylation of LC3 drive autophagy in response to D2-HG treatment *in vitro*.

**Figure 2.**
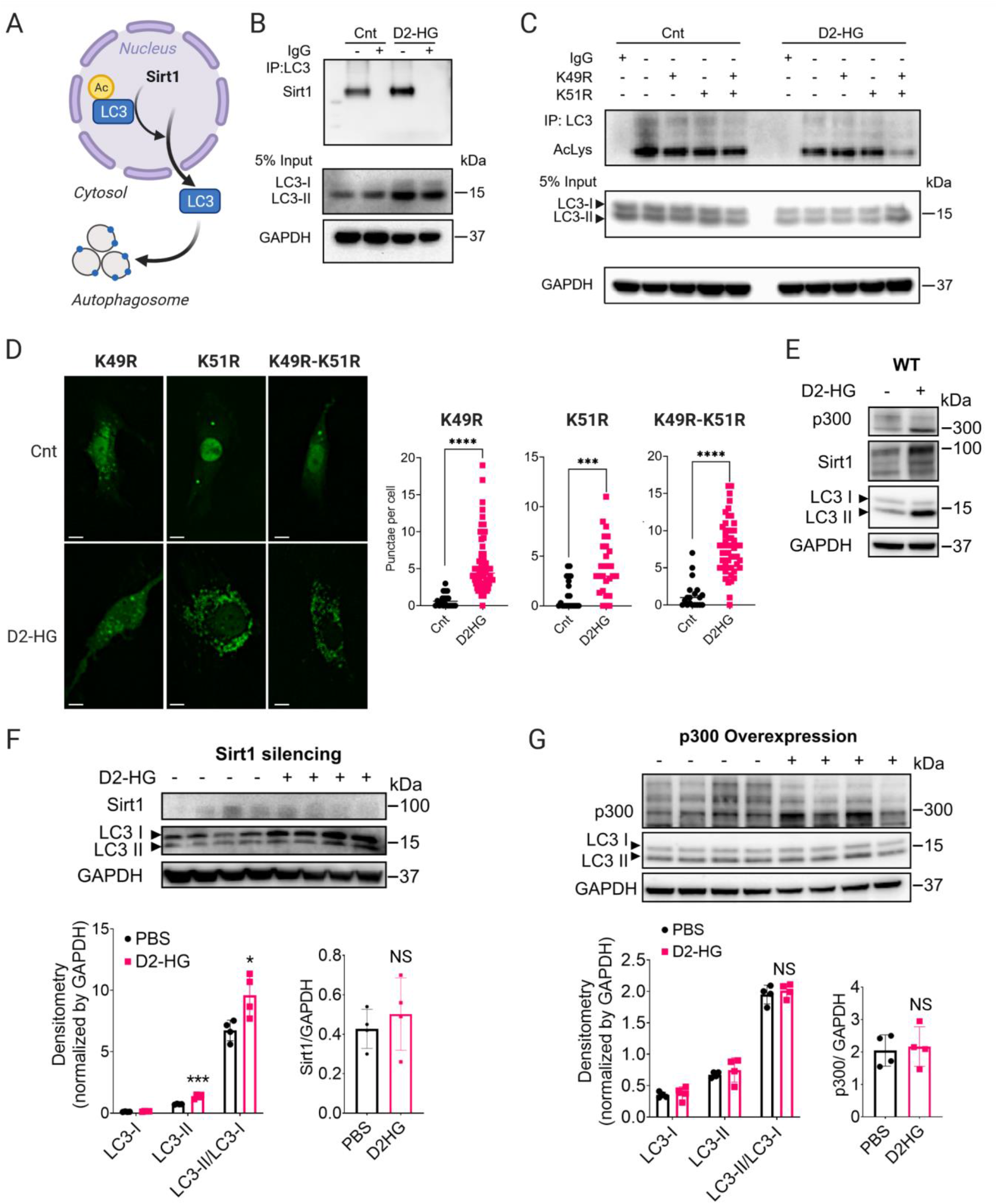
D2-HG-dependent deacetylation of LC3 is driven by Sirt1 in L6Ms. (**A**) Schematic of Sirt1-mediated deacetylation of LC3. (**B**) Representative western blotting depicting co-immunoprecipitation of LC3 and Sirt1 in differentiated L6Ms treated with or without D2-HG (1 mmol/L) for 24 h. Total expression of LC3-I, LC3-II, and GAPDH are depicted from 5% input of co-immunoprecipitation sample. Images are representative of n = 3 experiments. (**C**) LC3 was immunoprecipitated from L6 myotubes expressing wild type LC3 and mutant LC3 with a lysine to arginine replacement at position 49 and 51, respectively. The degree of acetylation was assessed using pan-acetyl-lysine antibody. Both wild type and mutant LC3 expressing cells were treated with or without D2-HG (1 mmol/L) for 24 h. (**D**) Representative live-cell images of differentiated L6Ms transfected with plasmid encoding a GFP-tagged LC3 (green) in wild type LC3 and mutant LC3 with a lysine to arginine replacement at position 49 and 51. Cells were treated with or without D2-HG (1 mmol/L) for 24 h, and puncta were counted for at least 100 cells per condition. Scale bars represent 10 μm. Data are mean ± s.d. \*\*\**P*-value<0.001, \*\*\*\**P*-value<0.0001. (**E**) Representative western blotting of Sirt1, p300, and LC3-I and LC3-II expression in L6 myotubes. (**F** and **G**) Silencing of Sirt1 expression (F) using siRNA and overexpression of p300 (G) reduces LC3-II level in L6 myotubes treated with D2-HG (*n* = 4 per group). Data are mean ± s.d. All *P*-values were determined by two-way analysis of variance (ANOVA).

### D2-HG promotes cachexia in vivo

To probe the biological importance and long-term consequences of D2-HG-mediated remodeling in skeletal muscle, wild type (WT) male and female mice (four animals per group; 10-week old) were treated for 30 days with either vehicle (PBS) or D2-HG (450 mg/kg body weight) through daily intraperitoneal injections (IP) (**Figure 3A**). We observed a reduction in *M. gastrocnemius* weight (both left and right muscles) in only male animals with D2-HG compared to placebo treatment after 30 days of treatment (**Figure 3B**). The total body weight was not significantly different between treated or untreated groups in both male and female animals (**Figure 3B**). Correspondingly, we found increased D2-HG levels in muscle tissue from male animals (**Figure 3C**). To assess frailty, we measured grip strength in mice by conducting a series of weekly *in vivo* weight-lifting tests during the 30-day treatment protocol (Contet *et al*, 2001; Deacon, 2013). Mice lifted weights ranging from 20 to 90 g for a maximum of 3 seconds (**Figure 3D**, see **Supplementary Methods** for details). In untreated control animals (male and female), grip strength increased in correlation with the animals’ overall growth (**Figure 3D**). Importantly, this growth-dependent increase in grip strength was significantly reduced in both males and females when treated with D2-HG for four weeks (male, *P*-value = 0.019; female, *P*-value = 0.0207; **Figure 3D**). Histological analysis of *M. gastrocnemius* sample revealed that the reduction in muscle mass in D2-HG treated male mice was caused by a reduction in the myofiber nuclei number (**Figure 3E**). Correspondingly, the LC3-II expression in *M. gastrocnemius* samples from male animals was reduced in D2-HG treated animals. These changes were not observed in female mice (Figure 3E). Total LC3 levels were almost depleted, with an accompanying 20-fold increase in p62 expression in skeletal muscle (**Figure 3F**). These findings show that prolonged treatment with D2-HG promotes disruption of LC3-mediated autophagy and p62 aggregation.

**Figure 3.**
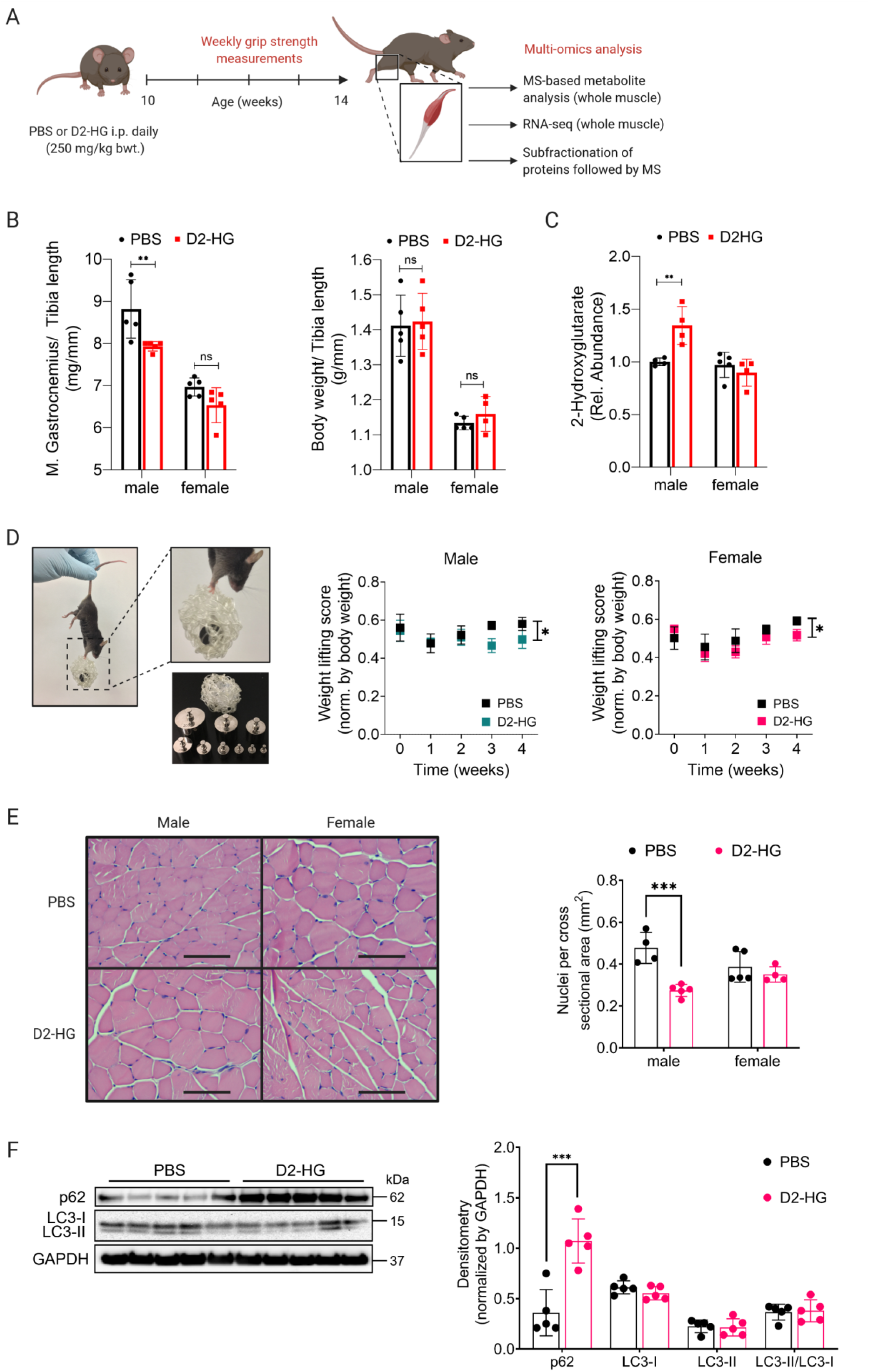
D2-HG promotes skeletal muscle atrophy *in vivo.* (**A**) Schematic for the experimental protocol. Whole skeletal muscle tissue from male and female mice were analyzed using RNA-sequencing (RNA-seq), mass spectrometry (MS) based targeted metabolomics, and MS-based untargeted proteomics using the IN-sequence workflow (see Methods for details). (Abbreviation: bwt, bodyweight). (**B**) Bodyweight and *M. gastrocnemius* weight normalized by tibia length of mice treated with PBS (control) and D2-HG (*n* = 4-5 mice per group and sex). Data are mean ± s.d. Statistical analysis by two-way analysis of variance (ANOVA). \*\**P*-value<0.01. (**C**) The relative abundance of 2-Hydroxyglutarate, normalized to the PBS-treated male and female group (*n* = 4-5 mice per group and sex). Data are mean ± s.d. \*\**P*-value<0.01. (**D**) Mice are lifting weights ranging from 20 g to 90 g. Normalized weight-lifting scores in male and female mice treated with or without D2-HG (250 mg/kg body weight) over the course of the experiment (4 weeks). Scores are representative of the grip strength in mice and normalized by body weight (*n* = 4-5 mice per group and sex). Data are mean ± s.d. ** *P*-value<0.01, \*\*\**P*-value<0.001. Statistical Analysis using a one unpaired t-test (two-step method by Benjamini, Krieger and Yekutieli) with a false discovery rate (FDR) of 5%. (**E**) Representative H&E images of *M. gastrocnemius* and quantification of nuclei per cross sectional areas from skeletal muscle sample in male and female mice treated with or without D2-HG (250 mg/kg body weight). scale bar = 100 μm; *n* = 4 per group. Data are mean ± s.d. ** *P*-value<0.01, \*\*\**P*-value<0.001. Statistical Analysis using a one unpaired t-test (two-step method by Benjamini, Krieger and Yekutieli) with a false discovery rate (FDR) of 5%. (**F**) Quantification of p62, LC3-I and LC3-II in skeletal muscle tissue from male mice (*n* = 5) treated with or without D2-HG (250 mg/kg body weight). Densitometry was normalized to the expression of GAPDH. Data are mean ± s.d. ** *P*-value<0.01, \*\*\**P*-value<0.001. Statistical Analysis using a one unpaired t-test (two-step method by Benjamini, Krieger and Yekutieli) with a false discovery rate (FDR) of 5%.

### Transcriptional and post-transcriptional regulation of oncometabolic stress

To investigate the extent of remodeling *in vivo* upon oncometabolic stress by D2-HG, we conducted an in-depth multi-omics analysis on skeletal muscle tissue sample (*M. gastrocnemius*) from mice with RNA sequencing (RNA-seq; transcriptomics) and LC-MS/MS for proteomics and metabolomics. Proteins were sub-fractionated into the highly abundant myofilamentous proteins, cytosolic proteins, and insoluble membrane proteins (Kane *et al*, 2007) prior to analysis by LC-MS/MS for identification and relative quantification (see **Supplementary Methods** for details). We identified a total of 46,079 transcripts using RNA-Seq, quantified 2,153 proteins from untargeted analysis using LC-MS/MS, and quantified 95 metabolites from targeted analysis using LC-MS/MS. PCA analysis of gene expression, as well as protein and metabolite abundances showed a clear separation between treatment and control groups (**Figure 4A**). In total, we identified 1,976 differentially expressed genes (FDR < 1%) and 170 differentially expressed proteins (FDR < 5%) that were enriched across 21 pathways using the Search Tool for the Retrieval of Interacting Genes/Proteins database (STRING, FDR < 0.01; **Figure 4B**) (Szklarczyk *et al*, 2019). Functional enrichment analysis showed that proteins in these clusters are part of cellular protein metabolic processes, regulation of chromatin assembly or disassembly, mitochondrial ATP provision, and muscle contraction (**Figure 4B**). Next, we reconstructed a protein-protein interaction (PPI) network of significantly regulated proteins using STRING (Szklarczyk *et al.*, 2019). The network consists of 70 proteins (nodes) and 170 protein-protein interactions (edges) (**Figure 4C**). D2-HG treatment differentially affected proteins that enriched in four main clusters: (1) mitochondrial respiratory chain complex assembly, (2) cell protein metabolic process, (3) cellular processes, and (4) chromatin assembly and disassembly. Normalized counts from RNA-seq could be matched with protein MS intensities for 1,346 genes independent of sex. Plotting individual RNA and protein ratios (D2-HG vs. PBS) revealed several proteins with differential expression at both the transcript and protein level (**Supplementary Figure 4A**). RNA and protein abundance ratios correlated only poorly (Pearson’s correlation coefficient, r = 8.9), indicating translational, miRNA or post-transcriptional regulation of many biological processes in response to oncometabolic stress in skeletal muscle. Intriguingly, several proteins that are part of autophagy regulation, lipid remodeling and known regulators of mitochondrial function, including the NADH dehydrogenase 1 alpha subcomplex subunit 13 (encoded by Ndufa 13), the quinone oxidoreductase (encoded by Cryz) and D-beta-hydroxybutyrate dehydrogenase (encoded by Bdh1), were only regulated at the protein level (**Supplementary Figure 4A**). Key regulators of chromatin organization and acetylation (Chd1 and histone 1.2) and autophagy regulation (Lamp1 and Bag3) showed increased protein expression in male and female mice treated with D2-HG (**Supplementary Figure 4B**). In contrast, gene and protein expression of Sirt2 and dynein were decreased for both male and female mice in treatment groups (**Supplementary Figure 4B**). These findings are consistent with chronic stress and epigenetic remodeling. Likewise, unsupervised hierarchical clustering of targeted metabolomics revealed four main clusters comprising 15 metabolic pathways using the Reactome pathway knowledgebase (Jassal *et al*, 2020) (**Supplementary Figure 5A**). Broadly, metabolites were enriched in pathways that are part of the pyrimidine metabolism, nucleotide metabolism, energy substrate metabolism, and DNA replication (**Supplementary Figure 5B**). Together these findings elucidate the adaptation to D2-HG treatment *in vivo* through the remodeling of chromatin to active, targeted gene programs, followed by changes in protein and metabolite abundances.

**Figure 4.**
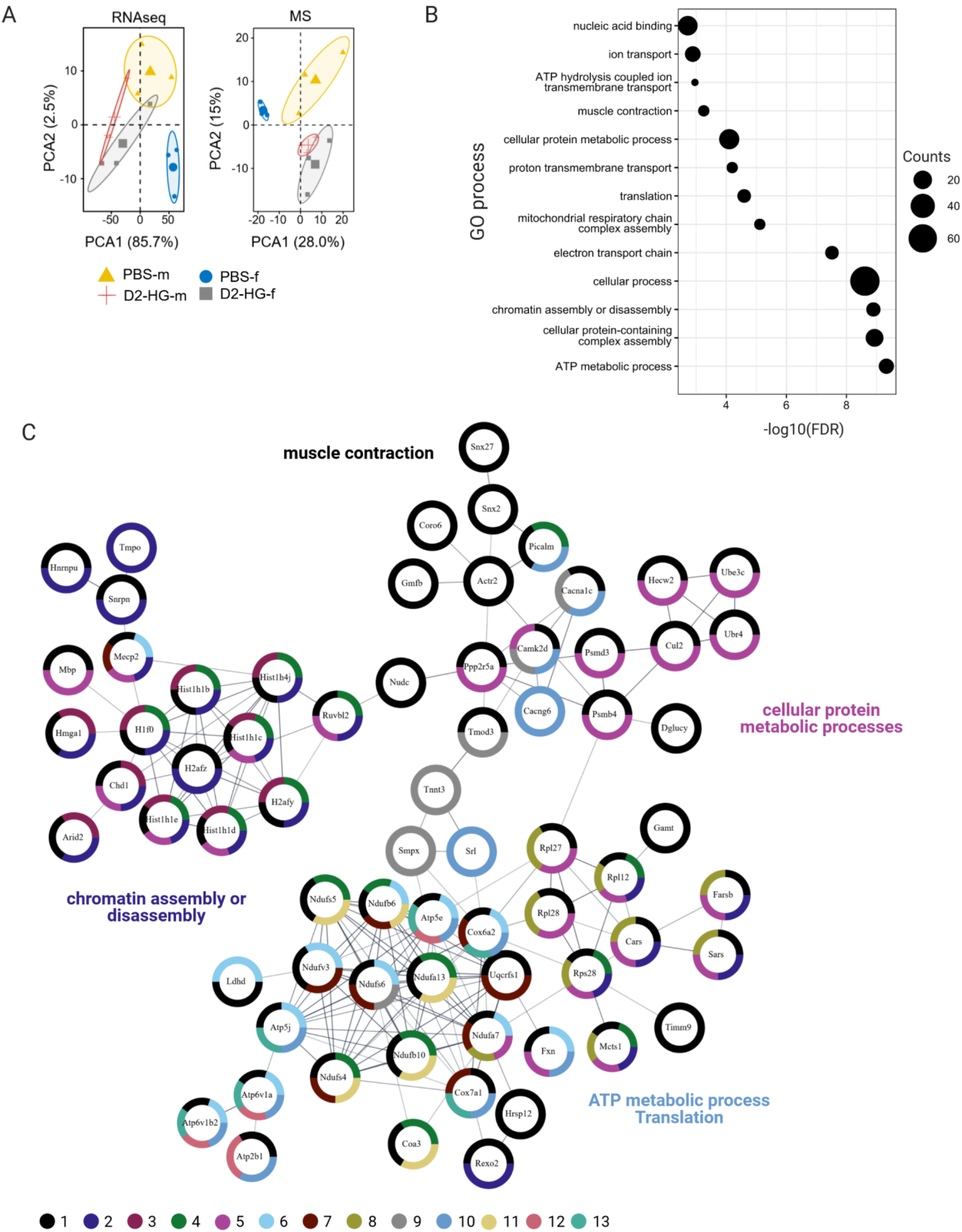
Combined transcriptomic and proteomic analysis of skeletal muscle remodeling upon oncometabolic stress. (**A**) PCA of RNA-sequencing (RNA-seq), MS-based proteomics, and MS-based metabolomics of skeletal muscle tissue from male (m) and female (f) mice treated with or without D2-HG (250 mg/kg body weight). Each data point represents a biological replicate. *n* = 4-5 mice per group and sex. Data is log2 normalized. (**B**) Enrichment analysis of significantly expressed proteins from MS-based proteomics using Gene Ontology (GO) annotations. The top-13 enriched terms after redundancy filtering were visualized according to negative log10 transformed FDR-values. The number of enriched genes is depicted by the size of each filled circle. (**C**) STRING protein-protein analysis of significantly expressed proteins from MS-based proteomics. Nodes represent proteins and edges represent protein-protein interactions. Functional analysis of clusters obtained by Markov clustering. GO annotation from (B) were visualized as split donut charts around the nodes as follows: 1 – muscle contraction; 2 – chromatin assembly or disassembly; 3 – nucleic acid binding; 4 – proton transmembrane transport; 5 cellular protein metabolic process; 6 – translation; 7 – mitochondrial respiratory chain complex assembly; 8 – electron transport; 9 – cellular protein-containing complex assembly; 10 – ATP metabolic process; 11 – ion transport; 12 – ATP metabolic process; 13 ATP hydrolysis coupled ion transmembrane transport. Proteins without any interaction partners within the network (singletons) are omitted from the visualization. (D-F) Comparison of gene and protein expression in skeletal muscle tissue for Chromodomain-helicase-DNA-binding-protein 1 (Chd1) and histone 1.2 (H1.2) (D), lysosome-associated membrane glycoprotein 1 (Lamp1) and Bcl-2-associated athano gene 3 (Bag3) (E), and Sirtuin 2 (Sirt 2) and Dynein (F). *n* = 4-5 mice per group and sex. Data are mean ± s.d. \**P*-value<0.05, \*\**P*-value<0.01, \*\*\**P*-value<0.001 and \*\*\*\**P*-value<0.0001. Statistical Analysis using a two-way ANOVA with a false discovery rate (FDR) of 5% for MS-proteomics and FDR of 1% for RNA-seq.

### Sex-dependent proteome and metabolome changes during cachexia

To compare the proteomic and transcriptomic datasets in terms of gene categories, we used a 2D annotation enrichment algorithm (Cox & Mann, 2012). This algorithm identifies both correlated and uncorrelated changes between two data dimensions. Male animals showed a reduction in gene expression and abundance of proteins that are part of oxidative phosphorylation and electron transport chain, whereas female animals upregulated the same set of proteins (**Figure 5A**). Likewise, there is notable metabolic remodeling that is both D2-HG-dependent and sex-dependent. Network analysis using MetaMapp (Barupal *et al.*, 2012) and cytoscape (Shannon *et al.*, 2003) revealed a sex-dependent metabolic profile of skeletal muscle adaptation in response to oncometabolite stress (**Supplementary Figure 6**). The metabolic profile in male mice treated with D2-HG showed increased relative abundance of pyruvate, lactate, erythrose 4-phosphate, α-KG, succinate, and amino acids (leucine, lysine, and methionine) (**Figure 5B**). In contrast, the abundance of fumarate, malate, and adenylsuccinate decreased in both sexes in response to D2-HG treatment (**Figure 5B**).

**Figure 5.**
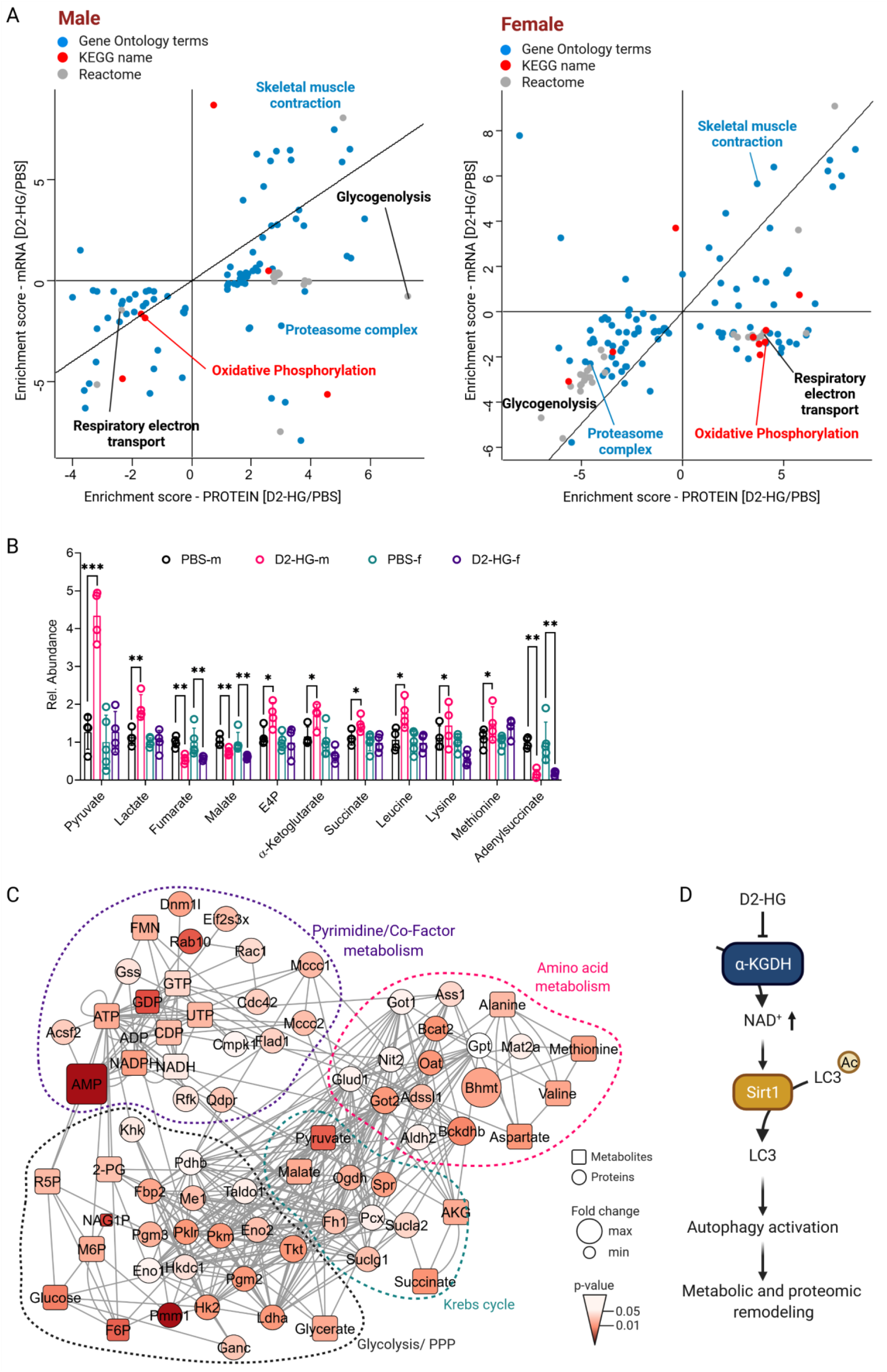
Sex-dependent adaptation in skeletal muscle during cachexia. (**A**) Two-dimensional annotation enrichment analysis based on transcriptome (RNA-sequencing) and proteome expression in skeletal muscle tissue from female and male mice treated with D2-HG for 30-days. Significant metabolic pathways (KEGG and Reactome) and gene ontology terms are distributed along the proteome and transcriptome change direction. Female and male mice show differentially up- or down-regulated metabolic pathways in response to D2-HG treatment *in vivo*. Types of databases used for annotations are color-coded as depicted in the legend. (**B**) Relative abundance of significantly up- or down-regulated metabolites in male and female mice treated with or without D2-HG. *n* = 4-5 mice per group and sex. Data are mean ± s.d. \**P*-value<0.05, \*\**P*-value<0.01, \*\*\**P*-value<0.001. Statistical Analysis using a two-way ANOVA with a false discovery rate (FDR) of 5% for MS-based metabolomics. (**C**) Network analysis of multi-omics data based on MS-based proteomics and targeted MS-based metabolomics. Nodes represent metabolites and proteins, while edges represent metabolite-metabolite, protein-protein, and metabolite-protein interactions. Metabolites and proteins are depicted as squares or circles, respectively. Nodes were color-coded by *P*-value and size represents the median fold change relative to untreated control animals. Statistical Analysis using a cut-off of 0.75 for Spearman’s rank correlation coefficient. (**D**) Summary of the main discoveries.

Next, we used the set of metabolites from targeted metabolomics to identify the directly interacting proteins (e.g., metabolic enzymes) using the Reactome pathway knowledgebase (Jassal *et al.*, 2020) and CardioNet (Karlstädt *et al*, 2012). In total we identified 361 identified protein-metabolite interactions (PMIs). The metabolites connected as a substrate, product, or co-factor to proteins in the initial network. The interaction between changes in metabolite level and protein expression was determined using a multiple linear regression (MLR) problem. We integrated 25 metabolites and 49 proteins into a network using a Spearman’s rank and Pearson correlation coefficient cut-off value of 0.75. The metabolic network is composed of 60 PMIs, 76 MMIs, and 229 PPIs (**Figure 5C**). We identified four main clusters of interactions that encompassed the (1) pyrimidine/co-factor metabolism, (2) amino acid metabolism, (3) Krebs cycle (or tricarboxylic acid cycle), as well as (4) glycolysis and the pentose phosphate pathway. Several core proteins interacted with multiple metabolites and PMIs preferentially occurred within a metabolic sub-network. For example, PMIs identified for glycolysis intermediates were both enriched in proteins for “glycolysis/pentose phosphate pathway,” and glucose-binging proteins were also enriched for the glycolysis (*P*-value<0.001). Our analysis shows the combined regulation of metabolic function through up- or down-regulation of proteins that are part of metabolic pathways and corresponding metabolite levels. For example, the upregulation of betaine-homocysteine S-methyltransferase (Bhmt) and branched-chain keto acid dehydrogenase E1 subunit beta (Bckdhb) expression correlated with an increased level of aspartate and methionine. Further, we identified five up-regulated proteins, Dnm1l, Rab10, Rac1, Cdc42 and Eif2s3x, that are part of cell signaling, organelle and microtubule organization, as well as myoblast fusion in mice and that are often co-expressed (Barabutis *et al*, 2018; Moore *et al*, 2019; Sylow *et al*, 2013; Vasyutina *et al*, 2009). The actin cytoskeleton-regulating GTPase, Rac1, is a known regulator of actin remodeling and contraction-induced glucose uptake through GLUT4 translocation. Correspondingly, we observed increased levels of AMP and glycolytic intermediates (e.g., glucose, fructose 6-phosphate), as well as an upregulation of proteins involved in glycolysis and the pentose phosphate pathway. Collectively, these results provide evidence that the oncometabolite D2-HG promotes skeletal muscle wasting through upregulation of autophagy (**Figure 5D**). Our multi-omics analysis further demonstrated the close interaction between protein and metabolite profiles in response to oncometabolic stress and expose metabolic vulnerabilities.

## Discussion

Our study revealed that the oncometabolite D2-HG promotes autophagy activation in skeletal muscle, and broad proteomic and metabolomic remodeling dependent on the sex. The response to oncometabolic stress includes early (e.g., energy substrate metabolism, altered redox state, and autophagy activation) and late events (e.g., structural protein remodeling, protein quality control, and chromatin remodeling). The combination of *in vitro* and *in vivo* multi-omics studies provided a comprehensive insight into the link between metabolic alteration and protein response pathways. We demonstrated that D2-HG impairs mitochondrial function in L6 myotubes, which increases NAD^+^ levels and activation of LC3-II, a key regulator of autophagy, through deacetylation by the nuclear deacetylase Sirt1. Consequently, increased expression of the protein acetylase p300 or decreased expression of Sirt1 prevented the activation of autophagy. In a mouse model of chronic D2-HG exposure, we also found that oncometabolic stress causes muscle atrophy, decreased grip strength, and increased expression of autophagy markers. Finally, a multi-omics analysis of transcriptomic, proteomics and metabolic data revealed a systems-wide sex-dependent remodeling in skeletal muscle in response to oncometabolic stress. Our analysis is a first step for the investigation of sex-dependent mechanisms in cachexia and oncometabolic stress, for instance, based on the enrichment for metabolic proteins.

The activation of proteolytic systems in mammalian cells is regulated by several pathways. Several lines of evidence indicate that D2-HG directly mediates autophagy and skeletal muscle atrophy. As there is no single autophagy marker, we used a series of experiments to answer whether D2-HG activates autophagy using cultured L6 myotubes. Gene expression shows a time-dependent activation of autophagy after treatment with D2-HG. Previous studies showed that LC3-II formation precedes an increased p62 expression during the activation of autophagy in nutrient starved cells (Gonzalez-Rodriguez *et al*, 2014; Pan *et al*, 2020; Runwal *et al*, 2019). Our data indicate that within 2 h after D2-HG treatment, Beclin1 expression increases rapidly, followed by a decreased p62 expression after 8 h. Correspondingly, we observed an increased LC3 expression within 12 h that was associated with increased Murf1 expression. These findings were corroborated in live-cell imaging using GFP-tagged LC3. Murf1 is an E3 ubiquitin ligase that mediates the ubiquitination and subsequent proteasomal degradation of muscle proteins (e.g., cardiac troponin I/TNNI3) and other sarcomere-associated proteins. Its role in muscle atrophy and hypertrophy is through regulating an anti-hypertrophic protein kinase C-mediated signaling pathway, which result in increased muscle protein degradation (Arya *et al*, 2004). Our data support the conclusion that D2-HG activates autophagy and protein degradation rapidly upon cellular exposure even in a nutrient-rich environment.

When using targeted metabolomics and functional mitochondria assays, we found that D2-HG impaired mitochondrial ATP provision and increased the NAD^+^ redox state. These findings agree with our previous studies in isolated working rat hearts, showing that D2-HG inhibits α-KGDH (Karlstaedt *et al.*, 2016). Several studies have shown an association between α-KG levels and autophagy. However, our findings indicate that the NAD^+^ redox state may be the primary driver of autophagy activation in myotubes. Using mutant LC3, we confirmed that autophagy activation in response to D2-HG is mediated through the deacetylation of LC3 by the nuclear deacetylase Sirt1. These findings are consistent with several recent reports that nuclear deacetylation of LC3 is driving autophagy during cell starvation (Huang *et al.*, 2015; Marino *et al*, 2014a; Shen *et al*, 2020; Yi *et al*, 2012; Zhao *et al*, 2016). Acetylation has emerged as an important post-translational modification that affects every step within the autophagic cascade. During periods of nutrient deficiency, cells initiate autophagy to replenish substrates for macromolecular synthesis. Our study advances this concept by providing evidence that oncometabolic reprogramming activates autophagy through a Sirt1-LC3 cascade even in a nutrient-rich environment. Metabolic stress evokes several cellular signaling pathways, including activation of AMPK and inactivation of mTOR. We did not observe an increased activation of AMPK in our *in vitro* experiments. The silencing of Sirt1 was sufficient to decrease autophagy, and p300 overexpression attenuated autophagy.

Together our data suggest that LC3-dependent mechanisms primarily drive autophagy activation in response to D2-HG. Our studies indicate that reductive α-KG metabolism is driving the protein acetylation changes within the autophagic cascade. Mechanistic challenges arise from linking metabolic changes to corresponding protein post-translational modifications and whether they are direct or indirect effects induced by metabolites. The arguments supporting our conclusions that autophagy is mediated directly through metabolic changes in the presence of D2-HG are the timing and concentration-dependent effects. As discussed above, metabolic alterations and acetylation of LC3 are sensitive to cellular α-KG levels and mitochondrial function. When D2-HG is elevated to 0.125 mM in L6 myotubes mitochondrial respiratory capacity is impaired, and both LC3 gene expression and lipidation increase within 12 h. Overexpression of p300 reverses the LC3 lipidation to control levels. Together these findings provide strong evidence that the observed effects are directly mediated through metabolic changes.

Using a multi-omics analysis of skeletal muscle from mice treated for one month with D2-HG, we identified a sex-specific metabolic, proteomic, and transcriptomic pattern that provides insight into the long-term consequences of cancer-induced remodeling. Specifically, we found that proteins involved in autophagy or proteasomal degradation machinery are increasingly expressed while structural proteins were decreased. Sirt1 has been found to drive autophagy via deacetylation and activation of FoxO3, a transcription factor that regulates the GTP-binding protein Rab7 and mediates autophagosome-lysosome fusion (Hariharan *et al*, 2010). Our analysis showed increased expression of lysosomal marker Lamp1 and FoxO3 at the protein and transcript levels. These findings suggest that initial metabolic response mechanisms that lead to activation of Sirt1 persist over time and induce a chronic activation of autophagy with broad metabolic and proteomic consequences.

Our data indicate that D2-HG treatment increases in both sexes the expression of proteins regulating chromatin organization, cytoskeleton organization, and cell signaling. These findings demonstrate the link between an initial metabolic response followed by targeted genome and proteome remodeling resulting in a new metabolic profile during prolonged adaptation. Further, our data show that the metabolic alterations induced by D2-HG affect the expression of proteins in a sex-dependent manner. Sex differences in muscle wasting and autophagy are increasingly recognized (Anderson *et al*, 2017; Miller *et al*, 2017; Montalvo *et al*, 2018; Yoon *et al*, 2018). Estrogens and androgens are important regulators of muscle mass and function (Spangenburg *et al*, 2012). Recent studies demonstrated that when autophagy is activated, acetylation of p300 is attenuated by Sirt1 or Sirt2 to modulate the degree of autophagic flux (Black *et al*, 2008; Han *et al*, 2008). Female mice are reported to have less basal ubiquitin-proteasome activity and greater autophagy activity compared to male mice (Cosper & Leinwand, 2011; Ogawa *et al*, 2015), while male mice appear to differentially regulate Sirt2 activity in response to ischemia (Shimizu *et al*, 2016). Together our data provide evidence for sex-dependent etiologies for cancer cachexia development and the need to optimize therapies for skeletal muscle pathologies based on biological sex. The role of sex in regulating autophagy and muscle wasting requires further studies.

Our findings shed new light on the mechanisms underlying the strong relationship between cancer and skeletal muscle loss by showing how oncometabolic stress activates autophagy. Our study points to a critical interplay between acetylation and deacetylation of proteins to regulate metabolic adaptation and proteome remodeling. It remains unclear how metabolic changes may impact epigenetic remodeling, and to what extent substrate replenishment can prevent the observed transcriptional and post-transcriptional changes. Our data clarify the importance of autophagy in cellular stress adaptation and provide insights into metabolic vulnerabilities driving skeletal muscle remodeling. Understanding the molecular mechanisms that enable muscle cells to adapt and compensate for oncometabolic stress will be necessary for therapeutic strategies targeting metabolic pathways in cachexia.

## Methods

### Animals

Animals were fed a standard laboratory chow, LabDiet 5001 (PMI Nutrition International, St. Louis, MO, USA). C57BL/6J Mice were obtained from Jackson Laboratory (Bar Harbor, ME, USA; CAT#000664, RRID: IMSR_JAX:000664), and both male and female mice were used in experiments.

### Cell lines, culture, and treatments

L6 myoblasts (rat skeletal muscle cell line) were purchased from American Type Culture Collection (ATCC, Manassas, VA, USA; ATCC CAT#CRL-1458, RRID: CVCL_0385). L6Ms were grown in Dulbecco’s Modified Eagle Medium (DMEM; Thermo Fisher Scientific, Hampton, NH, USA; CAT#10567022) supplemented with 10% (v/v) fetal bovine serum (FBS; Thermo Fisher Scientific, Hampton, NH, USA; CAT#16000044) and penicillin-streptomycin (100 units/mL; Thermo Fisher Scientific, Hampton, NH, USA; CAT#15140163) at 37°C and 5% CO2 in a humidified incubator. Cultures were grown for at least five passages before differentiation or transfection. L6Ms fuse in culture to form multinucleated myotubes and striated fibers. For differentiation into myotubes, L6Ms were grown in DMEM supplemented with 1% (v/v) FBS for 72 h to promote cell differentiation and fusion. Cells were then maintained at 37°C and 5% CO2 in a humidified incubator for the duration of the experiments, changing the media every 2-3 days. L6 myotubes were treated for 24 h with phosphate-buffered saline (PBS, Sigma Aldrich, St. Louis, MO, USA; CAT#506552), D-2-hydroxyglutarate (D2-HG, 0 to 1.0 mmol/L; Tocris Bioscience, Minneapolis, MN, USA; CAT#6124), or dimethyl 2-oxoglutarate (DMKG, 1 mmol/L, Sigma Aldrich, St. Louis, MO, USA; CAT#349631).

### Metabolic assays, RNA-sequencing, and mass spectrometry analysis

Details are provided in the expanded **Methods** section in the **Supplementary Information.**

### Data analysis and statistics

Statistical analysis was conducted using RStudio Desktop (v1.2.5042 for Linux, Boston, MA, USA) (RStudio Team, 2020), Bioconductor (Gentleman *et al*, 2004; Huber *et al*, 2015), and GraphPad Prism (version 8.1.2 for MacOS, GraphPad Software, La Jolla California USA, www.graphpad.com). Indicated sample sizes (n) represent individual tissue samples. For GFP-puncta counting, sample size (n) represents the number of cells analyzed from three or more independent experiments. Sample size and power calculations for *in vitro* and *in vivo* were based on Snedecor (Snedecor GW, 1989) and GPower (Faul, 2007) (version 3.1.9.2 for windows). The type 1 error and power were considered at 5% (*P-value* of 0.05) and 80%, respectively. The expected difference in the mean between groups was 50-30%, and the standard deviation of 25-12.5%. Further details are provided in the expanded **Methods** section in the **Supplementary Information**.

## Data and materials availability

The RNA-sequencing data have been deposited to the NCBI database (dataset identifier GSE159772). The MS proteomics data have been deposited to the ProteomeXchange Consortium via the PRIDE (Perez-Riverol *et al*, 2019) partner repository (dataset identifier PXD022137). The metabolomics data and networks files have been deposited to Mendeley.

## Acknowledgments

We thank the Histopathology Laboratory, Department of Pathology and Laboratory Medicine, at McGovern Medical School at The University of Texas Health Science Center at Houston for generating histology samples. Figures were created with BioRender.com. Funding: This work was supported by the American Heart Association (17POST33660221 to A.K.), the Burroughs Wellcome Fund (15RDM005 to A.K.) and National Institutes of Health (NIH) (R01-HL-061483 to H.T., K99-HL-141702 and R00-HL-141702 to A.K., R01-HL132075 and R01-HL144509 to R.A.G. and J.V.E. as co-PIs, K01-AI148593 to B.M.H.). The Metabolomics Core Facility at The University of Texas MD Anderson Cancer Center (P.L.L.) was supported by Cancer Prevention and Research Institute of Texas grant RP130397 and NIH grants S10OD012304-01, U01CA235510, and P30CA016672.

## Author Contributions

A.K. contributed to the conceptualization and project administration. A.K., J.V.E., B.H., P.L.L., R.G., H.T. contributed to the methodology. D.S., W.S, K.R. and J.V.E. (proteomics), A.K., L.T., and P.L.L. (metabolomics), A.K., A.D., A.G., B.H. (transcriptomics) contributed to omic data generation and/or processing. H.V., W.S., D.M., H.B.B. and S.A. contributed to the microscopy and histology data collection. A.K., H.V., B.D.G. contributed to *in vitro* cell culture studies; A.K., R.S., B.D.G. contributed to *in vivo* mouse studies; A.K., D.T., R.G. contributed to seahorse analysis. A.K. contributed to data curation. A.K. and H.V. contributed to data visualization. A.K., B.H., D.S., K.R., D.T., P.L.L. contributed to omic data analysis. A.K. wrote and prepared the original draft.

## References

Mendeley Data, doi: 10.17632/8843j9rfhk.1, http://dx.doi.org/10.17632/8843j9rfhk.1.

Amary MF, Bacsi K, Maggiani F, Damato S, Halai D, Berisha F, Pollock R, O’Donnell P, Grigoriadis A, Diss T et al (2011) IDH1 and IDH2 mutations are frequent events in central chondrosarcoma and central and periosteal chondromas but not in other mesenchymal tumours. J Pathol 224: 334–343

Anderson LJ, Liu H, Garcia JM (2017) Sex Differences in Muscle Wasting. Adv Exp Med Biol 1043: 153–197

Arya R, Kedar V, Hwang JR, McDonough H, Li HH, Taylor J, Patterson C (2004) Muscle ring finger protein-1 inhibits PKC{epsilon} activation and prevents cardiomyocyte hypertrophy. J Cell Biol 167: 1147–1159

Barabutis N, Dimitropoulou C, Gregory B, Catravas JD (2018) Wild-type p53 enhances endothelial barrier function by mediating RAC1 signalling and RhoA inhibition. J Cell Mol Med 22: 1792–1804

Baracco EE, Castoldi F, Durand S, Enot DP, Tadic J, Kainz K, Madeo F, Chery A, Izzo V, Maiuri MC et al (2019) alpha-Ketoglutarate inhibits autophagy. Aging (Albany NY) 11: 3418–3431

Barupal DK, Haldiya PK, Wohlgemuth G, Kind T, Kothari SL, Pinkerton KE, Fiehn O (2012) MetaMapp: mapping and visualizing metabolomic data by integrating information from biochemical pathways and chemical and mass spectral similarity. BMC Bioinformatics 13: 99

Black JC, Mosley A, Kitada T, Washburn M, Carey M (2008) The SIRT2 deacetylase regulates autoacetylation of p300. Mol Cell 32: 449–455

Bodine SC, Latres E, Baumhueter S, Lai VK, Nunez L, Clarke BA, Poueymirou WT, Panaro FJ, Na E, Dharmarajan K et al (2001) Identification of ubiquitin ligases required for skeletal muscle atrophy. Science 294: 1704–1708

Cancer Genome Atlas Research N, Ley TJ, Miller C, Ding L, Raphael BJ, Mungall AJ, Robertson A, Hoadley K, Triche TJ Jr., Laird PW et al (2013) Genomic and epigenomic landscapes of adult de novo acute myeloid leukemia. N Engl J Med 368: 2059–2074

Contet C, Rawlins JN, Deacon RM (2001) A comparison of 129S2/SvHsd and C57BL/6JOlaHsd mice on a test battery assessing sensorimotor, affective and cognitive behaviours: implications for the study of genetically modified mice. Behav Brain Res 124: 33–46

Cosper PF, Leinwand LA (2011) Cancer causes cardiac atrophy and autophagy in a sexually dimorphic manner. Cancer Res 71: 1710–1720

Cox J, Mann M (2012) 1D and 2D annotation enrichment: a statistical method integrating quantitative proteomics with complementary high-throughput data. BMC Bioinformatics 13 Suppl 16: S12

Deacon RM (2013) Measuring the strength of mice. J Vis Exp

DiNardo CD, Propert KJ, Loren AW, Paietta E, Sun Z, Levine RL, Straley KS, Yen K, Patel JP, Agresta S et al (2013) Serum 2-hydroxyglutarate levels predict isocitrate dehydrogenase mutations and clinical outcome in acute myeloid leukemia. Blood 121: 4917–4924

Evans WJ, Morley JE, Argiles J, Bales C, Baracos V, Guttridge D, Jatoi A, Kalantar-Zadeh K, Lochs H, Mantovani G et al (2008) Cachexia: a new definition. Clin Nutr 27: 793–799

Faul F, Erdfelder, E., Lang, A.-G., Buchner, A. (2007) G*Power 3: A flexible statistical power analysis program for the social, behavioral, and biomedical sciences. Behavior Research Methods 3: 175–191

Fearon K, Strasser F, Anker SD, Bosaeus I, Bruera E, Fainsinger RL, Jatoi A, Loprinzi C, MacDonald N, Mantovani G et al (2011) Definition and classification of cancer cachexia: an international consensus. Lancet Oncol 12: 489–495

Friesen DE, Baracos VE, Tuszynski JA (2015) Modeling the energetic cost of cancer as a result of altered energy metabolism: implications for cachexia. Theor Biol Med Model 12: 17

Gallagher IJ, Stephens NA, MacDonald AJ, Skipworth RJ, Husi H, Greig CA, Ross JA, Timmons JA, Fearon KC (2012) Suppression of skeletal muscle turnover in cancer cachexia: evidence from the transcriptome in sequential human muscle biopsies. Clin Cancer Res 18: 2817–2827

Gentleman RC, Carey VJ, Bates DM, Bolstad B, Dettling M, Dudoit S, Ellis B, Gautier L, Ge Y, Gentry J et al (2004) Bioconductor: open software development for computational biology and bioinformatics. Genome Biol 5: R80

Gomes MD, Lecker SH, Jagoe RT, Navon A, Goldberg AL (2001) Atrogin-1, a muscle-specific F-box protein highly expressed during muscle atrophy. Proc Natl Acad Sci U S A 98: 14440–14445

Gonzalez-Rodriguez A, Mayoral R, Agra N, Valdecantos MP, Pardo V, Miquilena-Colina ME, Vargas-Castrillon J, Lo Iacono O, Corazzari M, Fimia GM et al (2014) Impaired autophagic flux is associated with increased endoplasmic reticulum stress during the development of NAFLD. Cell Death Dis 5: e1179

Han Y, Jin YH, Kim YJ, Kang BY, Choi HJ, Kim DW, Yeo CY, Lee KY (2008) Acetylation of Sirt2 by p300 attenuates its deacetylase activity. Biochem Biophys Res Commun 375: 576–580

Hariharan N, Maejima Y, Nakae J, Paik J, Depinho RA, Sadoshima J (2010) Deacetylation of FoxO by Sirt1 Plays an Essential Role in Mediating Starvation-Induced Autophagy in Cardiac Myocytes. Circ Res 107: 1470–1482

Huang R, Xu Y, Wan W, Shou X, Qian J, You Z, Liu B, Chang C, Zhou T, Lippincott-Schwartz J et al (2015) Deacetylation of nuclear LC3 drives autophagy initiation under starvation. Mol Cell 57: 456–466

Huber W, Carey VJ, Gentleman R, Anders S, Carlson M, Carvalho BS, Bravo HC, Davis S, Gatto L, Girke T et al (2015) Orchestrating high-throughput genomic analysis with Bioconductor. Nat Methods 12: 115–121

Intlekofer AM, Dematteo RG, Venneti S, Finley LW, Lu C, Judkins AR, Rustenburg AS, Grinaway PB, Chodera JD, Cross JR et al (2015) Hypoxia Induces Production of L-2-Hydroxyglutarate. Cell Metab 22: 304–311

Intlekofer AM, Wang B, Liu H, Shah H, Carmona-Fontaine C, Rustenburg AS, Salah S, Gunner MR, Chodera JD, Cross JR et al (2017) L-2-Hydroxyglutarate production arises from noncanonical enzyme function at acidic pH. Nat Chem Biol 13: 494–500

Jassal B, Matthews L, Viteri G, Gong C, Lorente P, Fabregat A, Sidiropoulos K, Cook J, Gillespie M, Haw R et al (2020) The reactome pathway knowledgebase. Nucleic Acids Res 48: D498–D503

Kane LA, Neverova I, Van Eyk JE (2007) Subfractionation of heart tissue: the "in sequence" myofilament protein extraction of myocardial tissue. Methods Mol Biol 357: 87–90

Karlstädt A, Fliegner D, Kararigas G, Ruderisch HS, Regitz-Zagrosek V, Holzhütter H-G (2012) CardioNet: a human metabolic network suited for the study of cardiomyocyte metabolism. BMC Syst Biol 6: 114

Karlstaedt A, Zhang X, Vitrac H, Harmancey R, Vasquez H, Wang JH, Goodell MA, Taegtmeyer H (2016) Oncometabolite d-2-hydroxyglutarate impairs alpha-ketoglutarate dehydrogenase and contractile function in rodent heart. Proc Natl Acad Sci U S A 113: 10436–10441

Kir S, Spiegelman BM (2016) Cachexia & Brown Fat: A Burning Issue in Cancer. Trends Cancer 2: 461–463

Kranendijk M, Struys EA, Gibson KM, Wickenhagen WV, Abdenur JE, Buechner J, Christensen E, de Kremer RD, Errami A, Gissen P et al (2010a) Evidence for genetic heterogeneity in D-2-hydroxyglutaric aciduria. Hum Mutat 31: 279–283

Kranendijk M, Struys EA, Salomons GS, Van der Knaap MS, Jakobs C (2012) Progress in understanding 2-hydroxyglutaric acidurias. J Inherit Metab Dis 35: 571–587

Kranendijk M, Struys EA, van Schaftingen E, Gibson KM, Kanhai WA, van der Knaap MS, Amiel J, Buist NR, Das AM, de Klerk JB et al (2010b) IDH2 mutations in patients with D-2-hydroxyglutaric aciduria. Science 330: 336

Lee IH, Finkel T (2009) Regulation of autophagy by the p300 acetyltransferase. J Biol Chem 284: 6322–6328

Lok C (2015) Cachexia: The last illness. Nature 528: 182–183

Louis DN, Perry A, Reifenberger G, von Deimling A, Figarella-Branger D, Cavenee WK, Ohgaki H, Wiestler OD, Kleihues P, Ellison DW (2016) The 2016 World Health Organization Classification of Tumors of the Central Nervous System: a summary. Acta Neuropathol 131: 803–820

Lu C, Venneti S, Akalin A, Fang F, Ward PS, Dematteo RG, Intlekofer AM, Chen C, Ye J, Hameed M et al (2013) Induction of sarcomas by mutant IDH2. Genes Dev 27: 1986–1998

Marino G, Pietrocola F, Eisenberg T, Kong Y, Malik SA, Andryushkova A, Schroeder S, Pendl T, Harger A, Niso-Santano M et al (2014a) Regulation of autophagy by cytosolic acetyl-coenzyme A. Mol Cell 53: 710–725

Marino G, Pietrocola F, Kong Y, Eisenberg T, Hill JA, Madeo F, Kroemer G (2014b) Dimethyl alpha-ketoglutarate inhibits maladaptive autophagy in pressure overload-induced cardiomyopathy. Autophagy 10: 930–932

Miller AM, Shah RH, Pentsova EI, Pourmaleki M, Briggs S, Distefano N, Zheng Y, Skakodub A, Mehta SA, Campos C et al (2019) Tracking tumour evolution in glioma through liquid biopsies of cerebrospinal fluid. Nature 565: 654–658

Miller LR, Marks C, Becker JB, Hurn PD, Chen WJ, Woodruff T, McCarthy MM, Sohrabji F, Schiebinger L, Wetherington CL et al (2017) Considering sex as a biological variable in preclinical research. FASEB J 31: 29–34

Montalvo RN, Counts BR, Carson JA (2018) Understanding sex differences in the regulation of cancer-induced muscle wasting. Curr Opin Support Palliat Care 12: 394–403

Moore TM, Zhou Z, Cohn W, Norheim F, Lin AJ, Kalajian N, Strumwasser AR, Cory K, Whitney K, Ho T et al (2019) The impact of exercise on mitochondrial dynamics and the role of Drp1 in exercise performance and training adaptations in skeletal muscle. Mol Metab 21: 51–67

Nadtochiy SM, Urciuoli W, Zhang J, Schafer X, Munger J, Brookes PS (2015) Metabolomic profiling of the heart during acute ischemic preconditioning reveals a role for SIRT1 in rapid cardioprotective metabolic adaptation. J Mol Cell Cardiol 88: 64–72

Oberg AI, Dehvari N, Bengtsson T (2011) beta-Adrenergic inhibition of contractility in L6 skeletal muscle cells. PLoS One 6: e22304

Ogawa M, Kitakaze T, Harada N, Yamaji R (2015) Female-specific regulation of skeletal muscle mass by USP19 in young mice. J Endocrinol 225: 135–145

Pan B, Li J, Parajuli N, Tian Z, Wu P, Lewno MT, Zou J, Wang W, Bedford L, Mayer RJ et al (2020) The Calcineurin-TFEB-p62 Pathway Mediates the Activation of Cardiac Macroautophagy by Proteasomal Malfunction. Circ Res 127: 502–518

Pansuriya TC, van Eijk R, d’Adamo P, van Ruler MA, Kuijjer ML, Oosting J, Cleton-Jansen AM, van Oosterwijk JG, Verbeke SL, Meijer D et al (2011) Somatic mosaic IDH1 and IDH2 mutations are associated with enchondroma and spindle cell hemangioma in Ollier disease and Maffucci syndrome. Nat Genet 43: 1256–1261

Perez-Riverol Y, Csordas A, Bai J, Bernal-Llinares M, Hewapathirana S, Kundu DJ, Inuganti A, Griss J, Mayer G, Eisenacher M et al (2019) The PRIDE database and related tools and resources in 2019: improving support for quantification data. Nucleic Acids Res 47: D442–D450

RStudio Team, 2020. RStudio: Integrated Development for R. RStudio, PBC, Boston, MA

Runwal G, Stamatakou E, Siddiqi FH, Puri C, Zhu Y, Rubinsztein DC (2019) LC3-positive structures are prominent in autophagy-deficient cells. Sci Rep 9: 10147

Salazar-Degracia A, Granado-Martinez P, Millan-Sanchez A, Tang J, Pons-Carreto A, Barreiro E (2019) Reduced lung cancer burden by selective immunomodulators elicits improvements in muscle proteolysis and strength in cachectic mice. J Cell Physiol 234: 18041–18052

Shah A, Diculescu VC, Qureshi R, Oliveira-Brett AM (2010) Electrochemical reduction mechanism of camptothecin at glassy carbon electrode. Bioelectrochemistry 79: 173–178

Shannon P, Markiel A, Ozier O, Baliga NS, Wang JT, Ramage D, Amin N, Schwikowski B, Ideker T (2003) Cytoscape: a software environment for integrated models of biomolecular interaction networks. Genome Res 13: 2498–2504

Shen Q, Shi Y, Liu J, Su H, Huang J, Zhang Y, Peng C, Zhou T, Sun Q, Wan W et al (2020) Acetylation of STX17 (syntaxin 17) controls autophagosome maturation. Autophagy: 1–13

Shimizu K, Quillinan N, Orfila JE, Herson PS (2016) Sirtuin-2 mediates male specific neuronal injury following experimental cardiac arrest through activation of TRPM2 ion channels. Exp Neurol 275 Pt 1: 78–83

Smith KL, Tisdale MJ (1993) Increased protein degradation and decreased protein synthesis in skeletal muscle during cancer cachexia. Br J Cancer 67: 680–685

Snedecor GW CW (1989) Statistical methods. Iowa State University Press

Spangenburg EE, Geiger PC, Leinwand LA, Lowe DA (2012) Regulation of physiological and metabolic function of muscle by female sex steroids. Med Sci Sports Exerc 44: 1653–1662

Sylow L, Jensen TE, Kleinert M, Mouatt JR, Maarbjerg SJ, Jeppesen J, Prats C, Chiu TT, Boguslavsky S, Klip A et al (2013) Rac1 is a novel regulator of contraction-stimulated glucose uptake in skeletal muscle. Diabetes 62: 1139–1151

Szklarczyk D, Gable AL, Lyon D, Junge A, Wyder S, Huerta-Cepas J, Simonovic M, Doncheva NT, Morris JH, Bork P et al (2019) STRING v11: protein-protein association networks with increased coverage, supporting functional discovery in genome-wide experimental datasets. Nucleic Acids Res 47: D607–D613

VanderVeen BN, Fix DK, Carson JA (2017) Disrupted Skeletal Muscle Mitochondrial Dynamics, Mitophagy, and Biogenesis during Cancer Cachexia: A Role for Inflammation. Oxid Med Cell Longev 2017: 3292087

Vasyutina E, Martarelli B, Brakebusch C, Wende H, Birchmeier C (2009) The small G-proteins Rac1 and Cdc42 are essential for myoblast fusion in the mouse. Proc Natl Acad Sci U S A 106: 8935–8940

Yamamoto A, Tagawa Y, Yoshimori T, Moriyama Y, Masaki R, Tashiro Y (1998) Bafilomycin A1 prevents maturation of autophagic vacuoles by inhibiting fusion between autophagosomes and lysosomes in rat hepatoma cell line, H-4-II-E cells. Cell Struct Funct 23: 33–42

Yi C, Ma M, Ran L, Zheng J, Tong J, Zhu J, Ma C, Sun Y, Zhang S, Feng W et al (2012) Function and molecular mechanism of acetylation in autophagy regulation. Science 336: 474–477

Yoon SL, Grundmann O, Williams JJ, Gordan L, George TJ Jr., (2018) Body composition changes differ by gender in stomach, colorectal, and biliary cancer patients with cachexia: Results from a pilot study. Cancer Med 7: 3695–3703

Zhao S, Torres A, Henry RA, Trefely S, Wallace M, Lee JV, Carrer A, Sengupta A, Campbell SL, Kuo YM et al (2016) ATP-Citrate Lyase Controls a Glucose-to-Acetate Metabolic Switch. Cell Rep 17: 1037–1052

